# Maintaining arm control during self-triggered and unpredictable unloading perturbations

**DOI:** 10.1101/578450

**Authors:** Sasha Reschechtko, Anders S. Johansson, J. Andrew Pruszynski

**Affiliations:** Brain and Mind Institute; Robarts Research Institute; Western BrainsCAN; Department of Physiology and Pharmacology; Department of Psychology, Western University, London ON N6A 5B7 Canada; Department of Integrative Medical Biology, Physiology Section, Umeå University, Umeå Sweden

## Abstract

We often perform actions where we must break through some resistive force, but want to remain in control during this unpredictable transition; for example, when an object we are pushing on transitions from static to dynamic friction and begins to move. We designed a laboratory task to replicate this situation in which participants actively pushed against a robotic manipulandum until they exceeded an unpredictable threshold, at which point the manipulandum moved freely. Human participants were instructed to either stop the movement of the handle following this unloading perturbation, or to continue pushing. We found that participants were able to modulate their reflexes in response to this unpredictable and self-triggered unloading perturbation according to the instruction they were following, and that this reflex modulation could not be explained by pre-perturbation muscle state. However, in a second task, where participants reactively produced force during the pre-unloading phase in response to the robotic manipulandum to maintain a set hand position, they were unable to modulate their reflexes in the same task-dependent way. This occurred even though the forces they produced were matched to the first task and they had more time to prepare for the unloading event. We suggest this disparity occurs because of different neural circuits involved in posture and movement, meaning that participants in the first task did not require additional time to switch from postural to movement control.

## Introduction

To open a refrigerator door you have to pull on the handle with sufficient force to overcome the force sealing the door shut. The force sealing the door shut suddenly disappears once the seal breaks and the door begins to move. How you plan for and respond to this unpredictable transition between loaded and unloaded states depends on how you want to open the door. If you want to open the door just a little bit (i.e. to sneak a beer) you need to quickly dissipate the force you generated to break the seal because this force will subsequently act to swing the door open. On the other hand, if you intend to swing the door open completely (i.e. to put away your groceries) then dissipating this force is counterproductive.

The fridge door example represents a large class of agent-object interactions where the agent actively increases force to move the object and the force resisting movement unpredictably and suddenly decreases or disappears as a function of the force produced by the agent (e.g. almost every transition from static to dynamic friction). Here we investigate the behavioral strategies and underlying neural mechanisms associated with this class of interactions. We do so using a simple experimental paradigm where participants interact with two stiff walls, one behind and parallel to the other. Participants start by actively pushing against the first wall, which breaks at an unpredictable force level. When the first wall breaks, the arm is unloaded and begins to move. Participants are instructed to either break or not break the second wall.

Key to our design is its naturalism – motivated by our previous work in the jaw (Johansson 2014a,b) – where the participants are in active control of the force increase needed to trigger the sudden and unpredictable decrease in resistive force. This is contrast to previous studies along these lines where unloading was triggered by an external source (Angel et al. 1965, Asatryan & Feldman 1965, Crago et al. 1976, Dufossé et al. 1985, Paulignan et al. 1989, Latash & Gottlieb 1991, Lowrey et al. 2019). For example, in Angel and colleagues’ seminal study of reactions to unloading (1965) the participant counteracts a preset force level while controlling their position and passively waits for the unpredictable unloading event to be triggered by the experimenter cutting a wire. In contrast, in our paradigm unloading occurs while the participant voluntarily seeks the transition from loaded to unloaded state by actively increasing their applied force. Our approach is also in contrast with previous work focusing on self-triggered, and therefore predictable, unloading, as occurs when removing a load from one hand using the other hand (Lum et al. 1992), releasing a ball (Johansson & Westling 1988), or popping a balloon connected to a weight (Aruin & Latash 1995).

We report four principal findings. First, participants modulate the way they generate the force required to trigger the unloading perturbation depending on what they plan to do after unloading. Second, reflex responses are triggered approximately 40-60 ms following unloading and diverge within 100 ms after reflex onset depending on participants’ plans following unloading. Third, these reflex responses cannot be simply explained by pre-unloading muscle state; rather reflex processing appears to be actively modulated according to the instruction. Finally, the aforementioned reflex modulation requires that participants voluntarily generate force to trigger unloading, even though the unloading itself is unpredictable; when participants produce this force reactively, according to a prescribed trajectory enforced by the robot manipulandum, we find no reflex modulation.

## Methods

### Participants

A total of 22 participants (11 men, aged 20–36 yr.) participated in 1 of 2 experimental sessions. Twelve participants completed the first experimental session, which consisted of Experiment 1. Ten additional participants completed the second experimental session, which consisted of an abbreviated version of Experiment 1, as well as Experiments 2 and 3. In the second experimental session, the three Experiments were randomly interleaved. All participants were neurologically healthy and gave informed, written consent according to the declaration of Helsinki. The local ethical committee at Umeå University approved the study protocol. Experiments lasted approximately 2 hours, and participants were financially compensated for their time.

### Manipulandum

Experiments were conducted using a planar robotic manipulandum equipped with a vertical cylindrical handle (Endpoint KINARM; BKIN Technologies, Kingston ON, Canada; top panels of **Figure 1**). Participants grasped the handle to move the manipulandum in the horizontal plane against oppositional forces applied by the robot’s motors. Visual feedback regarding hand position, targets/obstacles and trial performance was displayed on a mirror halfway between the plane of the handle motion and a monitor positioned above the mirror. This configuration allows the cursor and targets to appear in the plane of the handle but occludes vision of the actual hand and manipulandum. The manipulandum recorded the position of the hand in the plane of motion and force transducers between the manipulandum and handle recorded the forces participants applied to the manipulandum. The robot was servo-controlled by a real-time computer at 4,000 Hz, while kinematic and kinetic data were digitally logged at 1,000 Hz on a separate computer.

**Figure 1.**
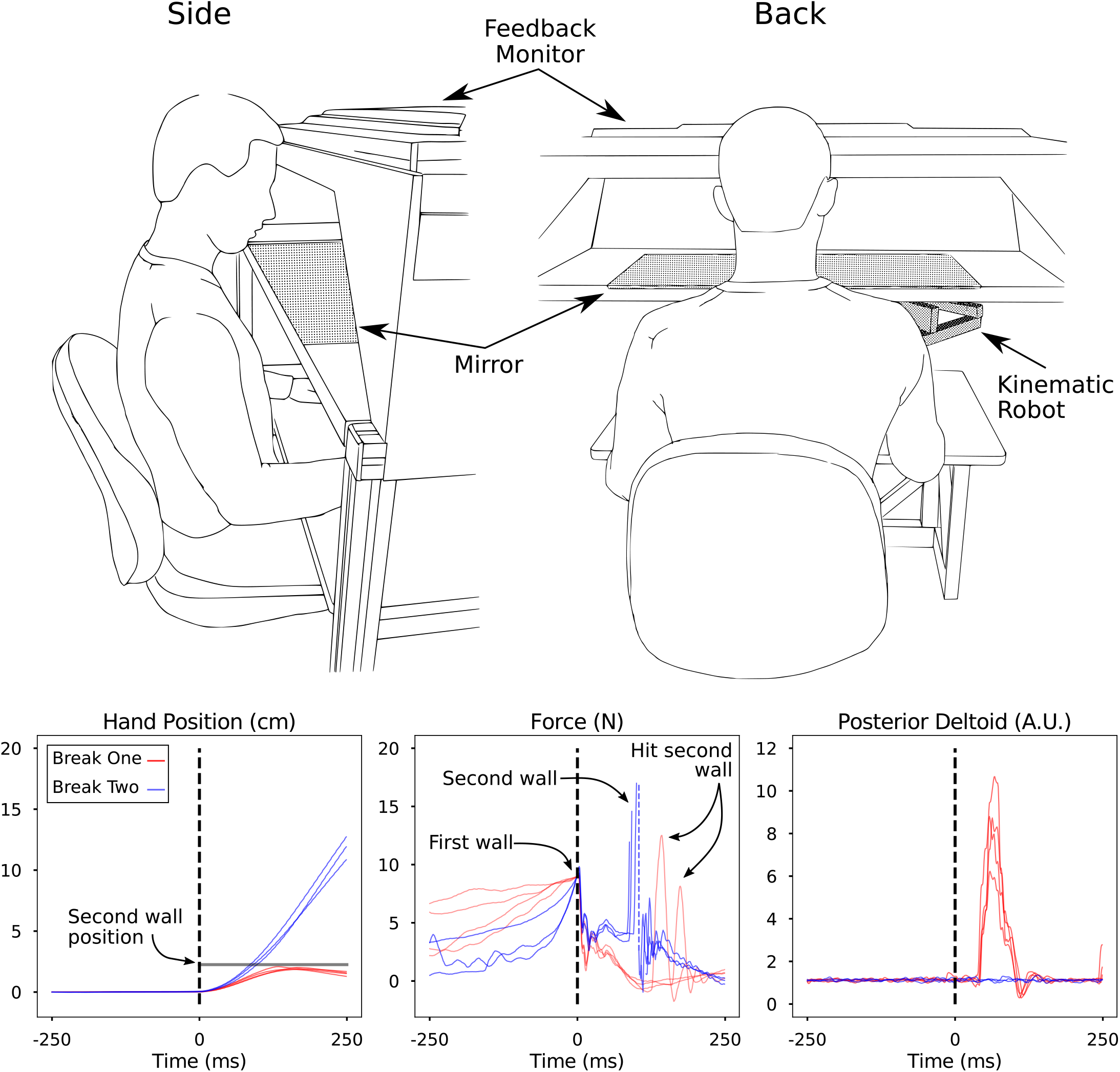
Upper Panels: Illustration of the experimental apparatus. Participants sat in front of a robot handle, which they grasped with their right hand. Hand position was hidden from view by a mirror, which reflected feedback displayed from a feedback monitor into the plane of the hand. Lower Panels: hand position, force applied to the robot handle, and posterior deltoid activation from an individual participant during one trial, centered around the unloading perturbation (dotted black line). Red traces refer to the trials when the participant was instructed to avoid breaking through the second wall (*Break One*), while blue traces are trials in which the participant was instructed to break through the second wall (*Break Two*). Dotted blue line in the force panel represents transient oscillations following breaking through the second wall, which were removed from the illustration for clarity. During some trials (annotated), the participant in the illustration hit the second wall with less force than was required to break through it (20N). Force traces in the *Break Two* condition do not reach 20N because, while the robot was servo controlled at 4kHz, the data presented were logged at 1kHz.

### Electromyography

Surface electromyograms (EMG) were obtained from three pairs of shoulder muscles: long head of the biceps, long head of the triceps, posterior deltoid, anterior deltoid, pectoralis major, and middle fibers of the trapezius. Prior to electrode placement, the skin was cleaned and abraded with rubbing alcohol and electrode contacts were covered with conductive gel. Electrodes (DE-2.1, Delsys, Boston, MA USA) were placed on the belly of the muscle and oriented parallel to the the muscle fibers. The quality of each EMG signal was assessed via a set of maneuvers known to elicit high levels of activation for each muscle in the horizontal plane of the task. EMG signals were recorded, amplified, and band-pass filtered (20–450 Hz) by a commercially available system (Bagnoli, Delsys) then digitally sampled at 1,000 Hz by the robotic manipulandum and synchronously with kinematic and kinetic data. For analysis, EMG data were normalized according to their mean activity during a task in which participants pushed the handle with a force of 20N, mimicking the configuration and force required in the rest of the experimental conditions. EMG amplitude was then quantified using an RMS envelope with a 10 ms moving average filter.

### Experimental Procedures

At the beginning of each experimental session, each participant sat comfortably in a chair holding the handle of the manipulandum with his or her right hand. The participants sat such that the upper arm was parallel to the torso with neutral shoulder rotation and the elbow flexed at ~90 degrees. The experimenter monitored participants’ arms during the experiment to ensure that participants did not abduct their shoulders; if abduction occurred, the trial was redone. Participants received visual feedback about hand position and the walls with which they interacted for the full duration of each trial, and at the end they were provided visual feedback about task performance (“GOOD” or “BAD” displayed on the screen).

Each experimental session began with a set of trials used to normalize EMG signals during off-line analysis. During these trials, subjects produced an isometric force of 20 N based visual feedback displayed on the screen in both push direction and pull directions. After these trials concluded, subjects performed 50 practice trials randomly selected from the protocol of Experiment 1 (these were discarded in the analysis) and subsequently participated experimental trials as described below.

#### Experiment 1a - Force dissipation and reflexes during unloading from a pushing movement

In Experiment 1a, participants were presented with two parallel walls which they could reach by moving the robotic handle away from the their body. Both of these walls were 8 cm long, 1.5 cm thick, and the walls were separated from one another by 2 cm. Participants were instructed to either *Break 1* or *Break 2;* in the *Break 1* condition, participants were required to produce enough force to break the closer wall but not break the second; in the *Break 2* condition, participants were required to break both walls. For the cohort of participants performing Experiment 1a during the first experimental session, the first wall broke at an unpredictable force level, which was varied between 5N and 23N in steps of 2N (for a total of 10 possible force levels), while the second wall always broke at 20N. For the cohort of participants performing Experiment 1a during the second experimental session, the first wall broke at a force level between 5N and 21N in steps of 4N, for a total of 5 possible force levels, while the second wall always broke at 20N.

Participants were given as much time as desired to break the first wall. *Break 1* trials were considered unsuccessful if subjects broke the second wall. *Break 2* trials were considered unsuccessful if the subject failed to break the second wall within 1500 ms after the first wall broke, or if the subject’s hand position deviated laterally so that it exited the space between the walls. Participants performed 4 successful repetitions of each force level for both the *Break 1* and *Break 2* instruction, with those in the first cohort performing a total of 80 successful trials in Experiment 1a and those in the second cohort performing 40 successful trials because they had fewer force levels for the first wall. The lower panels of **Figure 1** show one participant’s behavior during the 9N condition in terms of hand position, force produced against the manipulandum, and posterior deltoid muscle activation. As noted in the figure, participants sometimes “bounced” off the second wall in the *Break One* condition; this was considered a successful trial because they did not hit the second wall hard enough to break through it.

#### Experiment 1b - Force dissipation and reflexes during unloading from a pulling movement

The participants in Experiment 1a also performed the same procedure outlined above with a pulling action. The location of the virtual wall was moved to the other side of the hand’s start position, so that a pulling action was required to break the instructed number of walls.

#### Experiment 2 - Alteration of inter-wall distance

In Experiment 2, the distance between walls was altered. In half of the trials it was 2 cm (as in Experiment 1), while in the other half it was 6 cm. For these trials, the force level at which the first wall broke was varied between 5N and 21N in steps of 4N for a total of 5 levels. Participants performed 4 successful repetitions of each force level, inter-wall distance, instruction combination, for a total of 80 trials.

#### Experiment 3 - Passive force increase

In Experiment 3, we investigated whether reflex responses were modulated when participants could not control the force profile before they broke through the first wall. Participants were instructed to keep the robotic handle in a set position (circular target, 1.0 cm in diameter at same location where they initiated wall breaking in Experiments 1 and 2) while the manipulandum exerted an increasing force. In some trials, participants were instructed to stay in the initial position after the force was suddenly turned off by the robot (similar to Break One tasks). In other trials, subjects were instructed to respond to the release by pushing the handle away from their body to reach a new final location (similar to Break Two tasks). The force profile which participants needed to counter was varied across trials, a low force rate (10 N/s) and a high force rate (40 N/s). The force applied to the participant was released unpredictably at levels between 5N and 21N in steps of 4N for a total of 5 levels. Participants performed 4 successful repetitions of each first wall force level/force rate/instruction combination for a total of 80 successful trials.

### Data analysis

Data were processed offline using MATLAB (The Mathworks, Natick, MA, USA) to extract data from the data structures output by the kinematic robot and Python 3 (Python Software Foundation, Wilmington, DE, USA) for subsequent analysis. For each trial, a 1 s analysis epoch was defined for all data collected. These epochs were aligned such that their midpoints were the first wall break in Experiments 1 and 2, and the time of force release in Experiment 3. Subsequent analysis was carried out on these aligned data to recover several variables of interest, as described below.

#### Force Rate

We assessed force rate using force data collected at the handle of the manipulandum. We analyzed mean force rate for each break force in each condition over a 50 ms window which ended at the unloading event (first wall break). We chose this short window because the epoch of force generation was highly variable between levels of break force and instruction conditions.

#### EMG Preceding Unloading

We analyzed EMG leading up to the unloading event over the same window was force rate. We analyzed two quantities reflecting whole-arm EMG activity: generalized agonist and antagonist activation, defined as the sum of normalized EMG activity in all three agonist or antagonist muscles.

#### Reflex Onset and Magnitude

We assessed reflex onset via EMG. In each trial, EMG data for each muscle were median filtered; we then extracted the mean value and standard deviation for the “silent period” observed during the first 20 ms following unloading. We identified the reflex onset for a given muscle as the time following unloading when a given muscle’s activity exceeded 3 standard deviations above the mean EMG value during the silent period. Reflex magnitude was subsequently defined for each trial with identifiable reflexes as the average EMG amplitude between the time of onset and 100ms following the reflex onset.

#### Reflex Gain

We investigated whether reflex magnitudes in the posterior deltoid following the unloading perturbation were related the activity of the posterior deltoid preceding unloading by using linear regression analysis. We z-scored reflex activity and pre-unloading activity separately, and then regressed reflex activity on pre-unloading activity. The slope in this analysis represents the amount that reflex activity increases with increasing baseline (pre-perturbation) activity, which is expected due to so called “automatic gain scaling” (Bedingham & Tatton 1984, Matthews 1986; Pruszynski et al., 2009). The intercept of the regression represents other potential sources of reflex modulation that are not consistent with classical gain scaling.

### Statistics

To determine the time at which reflex responses under each instruction diverged from each other, we ran independent sample two-sided t-tests on each point in the timeseries of posterior deltoid EMG activity for all trials under each condition (four samples per condition at each time point) following the unloading event. We define the point of divergence for an individual participant and level of wall force as the time point at which three consecutive t-tests were significant. Before running the t-tests, EMG activity at the time of unloading was subtracted from each trial to ensure the difference detected did not result from the level of activity at the time of unloading. Note that this results in divergence times reported being later than visual inspection would indicate. T-tests were run using *ttest_ind* command in the *Scipy* library for Python (Jones et al. 2001).

Repeated-measures ANOVAs were carried out using R (R Core Team 2018) with the *lme4* (Bates et al. 2015), *limerTest* (Kuznetsova et al 2014), *pbkrtest* (Halekoh & Højsgaard 2014) and *emmeans* (Lenth 2018) packages. Repeated-measures ANOVAs were carried out using linear mixed model analysis (*lme4*) with independent intercepts and individual subjects treated as random variables and reduced maximum likelihood fitting. P-values for effects were obtained using the Kenward-Roger (Kenward & Roger 1997) approximation for denominator degrees of freedom (implemented through *limerTest* and *pbkrtest*). Luke (2017) indicates this approach for estimating p-values is well-calibrated for small sample sizes. Code for these analyses is adapted from Winter (2013).

## Results

### Experiment 1a

This experiment investigated how participants prepare for and respond to unpredictable self-triggered unloading perturbations depending on what they are instructed to do after the perturbation. In agreement with previous work studying similar perturbations in the jaw (Johansson et al. 2014a,b), we observed instruction-dependent differences both before and after the unloading perturbation.

All participants (n = 22) modified their behavior before the unloading event depending on which instruction they were given (**Figure 2**; exemplar participant in lower panels of **Figure 1**). As they produced force against the robot handle to break through the first virtual wall, the rate at which they increased this force was higher and depended on the strength of the wall under the “Break Two” instruction but not under the Break One instruction (*Break Force × Instruction: F_4,189_* = 10.876; *P* = 5.88e^−8^). Across break force levels, average force rate under the “Break One” instruction was 50.49 ± 6.8 N/s, while it was 119.41± 7.47 N/s under the “Break Two” instruction. Agonist and antagonist EMG activity showed modulation under the Break One instruction but not under the Break Two instruction (Agonist *Break Force × Instruction: F_4,189_* = 5.14; *P* = 5.91e^−4^; Antagonist: *F*_4,189_ = 4.30; *P* = 0.0024) such that agonist and antagonist activation increased with the strength of the wall for Break One but was relatively unchanged for Break Two, even though force rate is higher for Break Two. That is, participants only increased co-contraction when they needed to decelerate from higher force levels, rather than always producing high co-contraction with high force levels.

**Figure 2.**
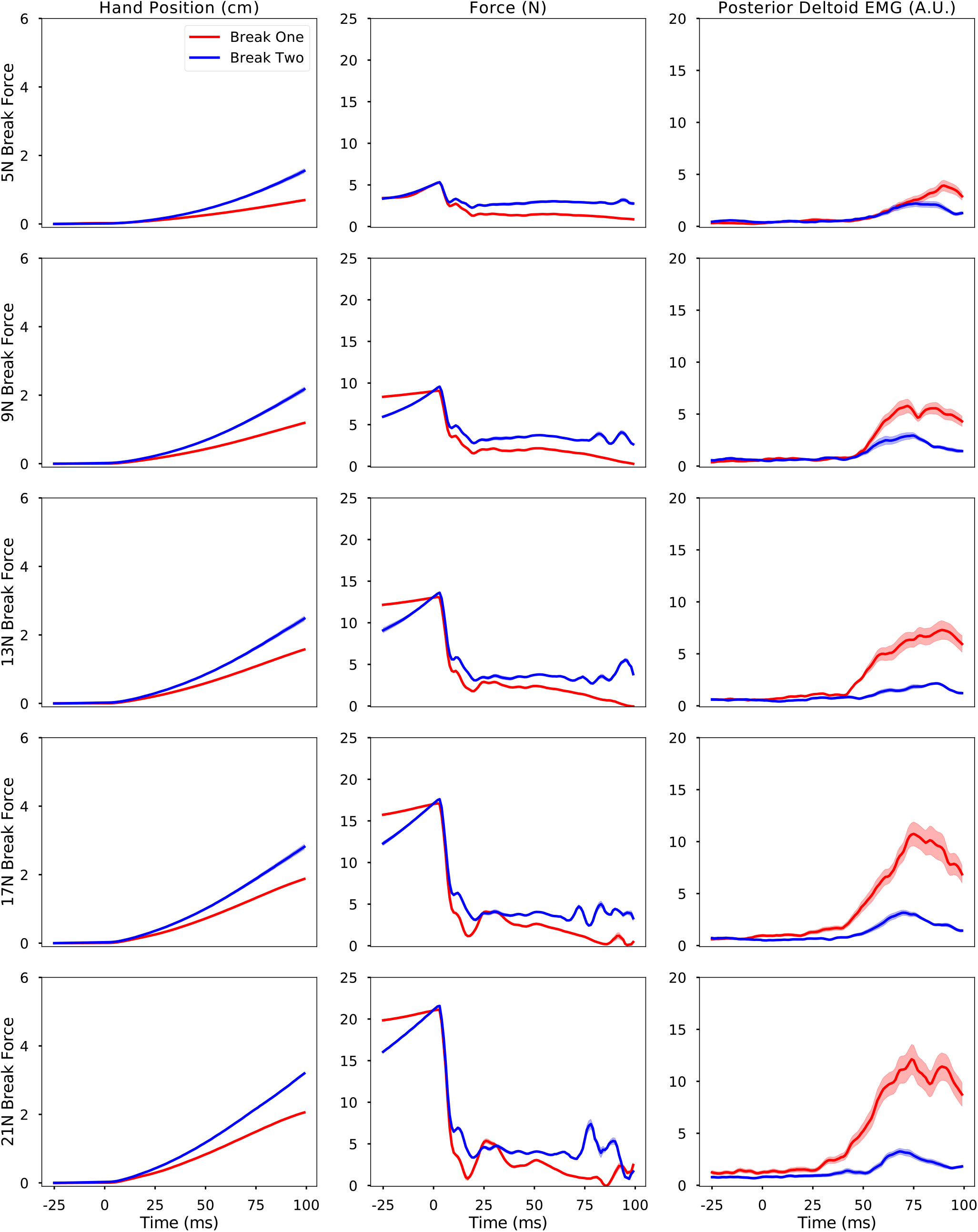
Thin lines represent individual participant (n = 22) mean performance during Experiment 1, aligned with perturbation onset at time = 0. Red traces refer to trials when participants were instructed to stop before breaking the second wall following the perturbation; blue traces are trials in which participants were instructed to continue to push through the second wall following the perturbation. Thick traces represent across-subject means. Rows refer to the force level at which the perturbation was triggered. Column 1: kinematics of the hand perpendicular to the wall (distance in cm); Column 2: force applied to the robot handle in the direction of the wall (force in N); Column 3: EMG activity from the posterior deltoid, a primary antagonist (normalized by activity when applying 20N in the direction of the wall).

Following the unloading perturbation, measures of hand position, force exerted on the robot handle, and EMG in the posterior deltoid (the primary antagonist muscle) all diverged rapidly depending on the instruction (**Figure 3**). We quantified the time of divergence for each variable using sequential t-tests (see Methods; **Figure 3**, top row). Across levels of Break Force, both hand position and posterior deltoid EMG were significantly different under Break One and Break Two instructions for approximately 80% of subjects by 100ms following unloading, while force applied to the robot handle was different for approximately 95% of subjects by 100ms following unloading.

**Figure 3.**
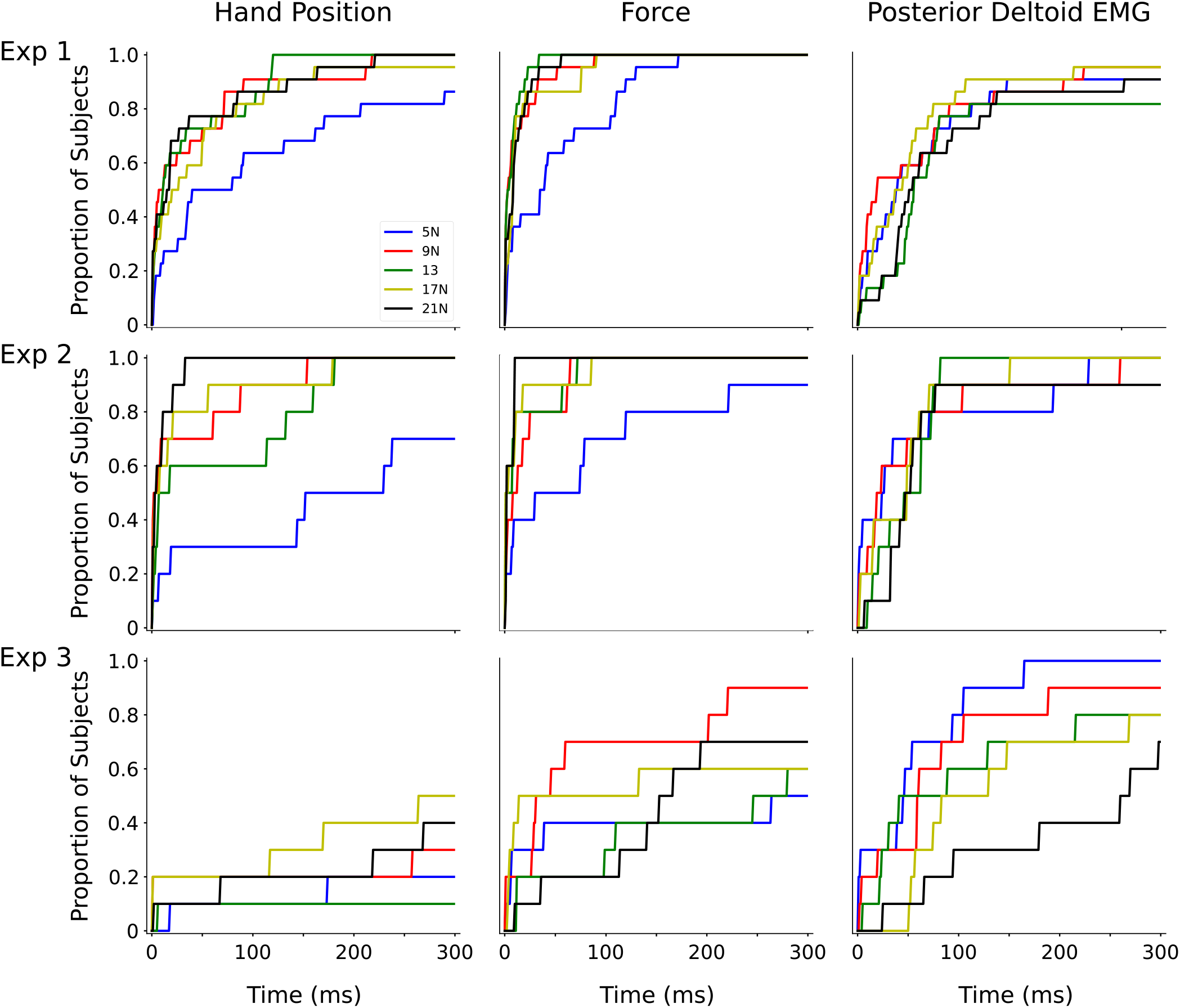
Proportion of participants showing differentiated responses during sequential t-tests comparing *Break One* and *Break Two* behaviors at a given time following the perturbation. Columns refer to the measure tested, while rows refer to the experiment number.

We also identified reflex onsets in the posterior deltoid (see Methods). On average, we found that the onset was earlier under “Break One” (41.16 ± 3.20 ms; mean ± SEM) than “Break Two” (53.26 ± 6.69 ms). To ascertain whether reflexes were modulated according to instruction, we compared the mean value of posterior deltoid EMG between reflex onset and 100 ms following perturbation onset. We found that EMG in the posterior deltoid was larger under the Break One instruction, where its activity would serve to decelerate the hand, than under the Break Two instruction (where its activity would be counterproductive), and that it scaled with break force under the Break One instruction only (*Break Force × Instruction* i nteraction: *F*_4,186_ = 4.75; *P* = 0.0011). ‘

Because we previously found that EMG in the posterior deltoid was higher before perturbation onset, we used linear regression to analyze the relationship between the magnitude of the reflex response and its EMG level during 50 ms preceding the unloading event of the first cohort (n = 12) of participants, who experienced more levels of break force. If differences in posterior deltoid reflex magnitude were completely determined by the loaded muscle state (as might be expected given reflex gain-scaling, cf. Matthews 1986), we would expect regression coefficients and intercepts to be identical. However, we found that the relationship between posterior deltoid activation and reflex magnitude was different depending on the instruction given to the subject. We found that, across the population, participants show no reliable modulation of the gain of this relationship (slope) according to instruction (paired t-test: *T*_11_ = 0.84; *P* = 0.416), whereas the intercept of the response function was reliably different such that a given level of posterior deltoid activation was associated with a larger reflex response under the “Break One” instruction (*T*_11_ = 5.24; *P* = 2.77e^−4^). These results are illustrated in the first panel of Figure 3.

### Experiment 1b

We carried out experiment 1b to assess the generality of the responses observed in experiment 1a. We found qualitatively similar results to all those reported for Experiment 1a, except that agonists and antagonists were reversed. Most importantly, task-dependent reflex modulation was evident in pectoralis (which is the antagonist here): regressing reflex magnitude on posterior deltoid activation preceding unloading, we again found a significant change in intercept (*T*_11_ = 7.82; *P* = 8.07e-6) depending on instruction, but not on slope (*T*_11_ = −1.11; *P* = 0.29.

### Experiment 2

Experiment 1 established that participants prepare for a self-triggered unloading perturbation differently depending on how they were instructed to respond to this perturbation. In Experiment 2, we investigated whether these task-dependent strategies continued to differ when the task was less difficult. In a second cohort of participants (n = 10), we decreased the difficulty of the task by increasing the distance between the walls, thereby giving participants more distance to decelerate their arm after the unloading perturbation, with the expectation that task-dependent modulation would decrease when the task was easier.

As in Experiment 1, participants’ behaviors leading up to the unloading perturbation differed according to the instruction they were given (**Figure 5**). Even when the second wall was farther away, we continued to observe modulation of generalized agonist and antagonist musculature under the Break One instruction (*Break Force × Instruction* interaction in the 3-way ANOVA *Break Force × Instruction × Wall Distance*; agonist: *F*_4,171_ = 3.75; *P* = 0.006, antagonist: *F*_4,171_ = 3.25; *P* = 0.013) before participants incurred the unloading perturbation. This relationship was not significantly affected by the distance to the second wall.

Participants’ reflex responses to the unloading perturbation also diverged quickly when the wall was farther away (**Figure 5**). Posterior deltoid reflex magnitudes were modulated with break force for Break One but not Break Two (*Break Force × Instruction* interaction in the 3-way ANOVA *Break Force × Instruction × Wall Distance F*_4,167_ = 3.61; *P* = 0.0075), and this modulation was not significantly affected by the distance to the second wall. On average, participants’ reflex responses to the perturbation under the Break One instruction (near: 51.21 ± 3.73 ms; far: 45.83 ± 3.05 ms) were earlier than under the “Break Two” instruction (near: 68.62 ± 9.88 ms; far: 79.43 ± 11.0 ms). By 100 ms following the reflex onset, position was significantly different under Break One and Break Two instructions for approximately 67% of participants, while posterior deltoid EMG was significantly different for 70% of participants and force applied to the robot handle was different for 90% of participants (as assessed by sequential t-tests). Regression of posterior deltoid reflex activity on the pre-loading baseline also indicate similar behaviors regardless of the distance between the two walls: in both conditions, the intercept was reliably different between instructions (near wall: *T*_9_ = 3.15; *P* = 0.012, far wall: *T*_9_ = 5.77; *P* = 2.7e^−4^) while the slope was not reliably different for either wall distance (near wall: *T*_9_ = 0.41; *P* = 0.69, far wall: *T*_9_ = 0.39; *P* = 0.70).

### Experiment 3

In Experiment 3, we focused on the role of self-triggering the unloading perturbation. The same cohort of participants who participated in Experiment 2 also performed a task where they were instructed to maintain their hand in a specific position (equivalent to the position they were at when pushing against the first wall in Experiments 1 and 2) as the robotic handle smoothly ramped up force until it exceeded one of the force thresholds from Experiment 2. On different trials, participants were given one of two different instructions: in one, they were to stay in the same position after the oppositional force was unloaded (similar to Break One); in the second, they were to continue to push in the same direction they had been pushing such that they moved their hand in that direction towards a new final position (similar to Break Two).

Because force production in Experiment 3 was set by the robot, force rates preceding unloading did not change depending on the instruction to move after unloading. EMG activity preceding unloading did not show modulation according to the instruction given to subjects, but it did vary somewhat depending on the break force and rate at which the robot increased force (*Break Force × Force Rate* interaction for agonist: *F*_4,171_ = 2.72; *P* = 0.048, antagonist: *F*_4,171_ = 4.44; *P* = 0.0019).

Compared to Experiments 1 and 2, behaviors took longer to diverge following the unloading event (**Figure 6**). Hand position showed minimal modulation according to instruction over the first 100 ms following unloading, while force applied to the robot handle and posterior deltoid activity also showed slower modulation than in Experiments 1 and 2 (bottom row in **Figure 3**), with only 20-30% of subjects showing reliable task-dependent behaviors during the first 100 ms across force levels. Similarly, reflex magnitudes in posterior deltoid were not modulated according to instruction (Main effect of *Instruction* i n 3-way ANOVA *Break Force × Instruction × Force Rate: F*_1,171_ = 0.28; *P* = 0.6, without significant interactions). Finally, in contrast to that seen in Experiments 1 and 2, the relationship between reflex magnitude in the posterior deltoid and pre-unloading activation in posterior deltoid did not display reliable modulation related to movement instruction (Slope: *T*_9_ = −0.07 *P* = 0.94; *T*_9_ = −0.24 Intercept: *P* = 0.81), as shown in the third panel of **Figure 4**.

**Figure 4.**
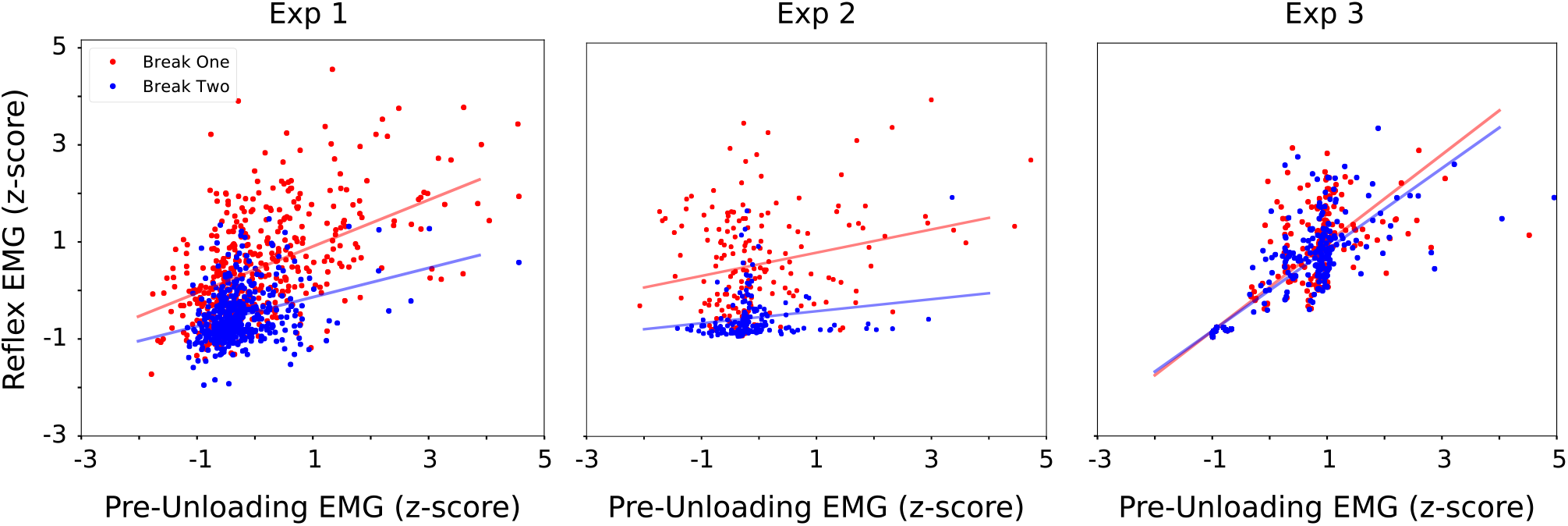
Regression of posterior deltoid reflex magnitude on the state preceding the unloading perturbation. Slopes of the regression correspond to automatic gain scaling, while vertical displacement indicates a task-specific alterations in response independent of gain scaling. Experiment 2 data are plotted for the task when the second wall was in the “far” position, and Experiment 3 data are plotted for the “fast” force ramp condition.

**Figure 5.**
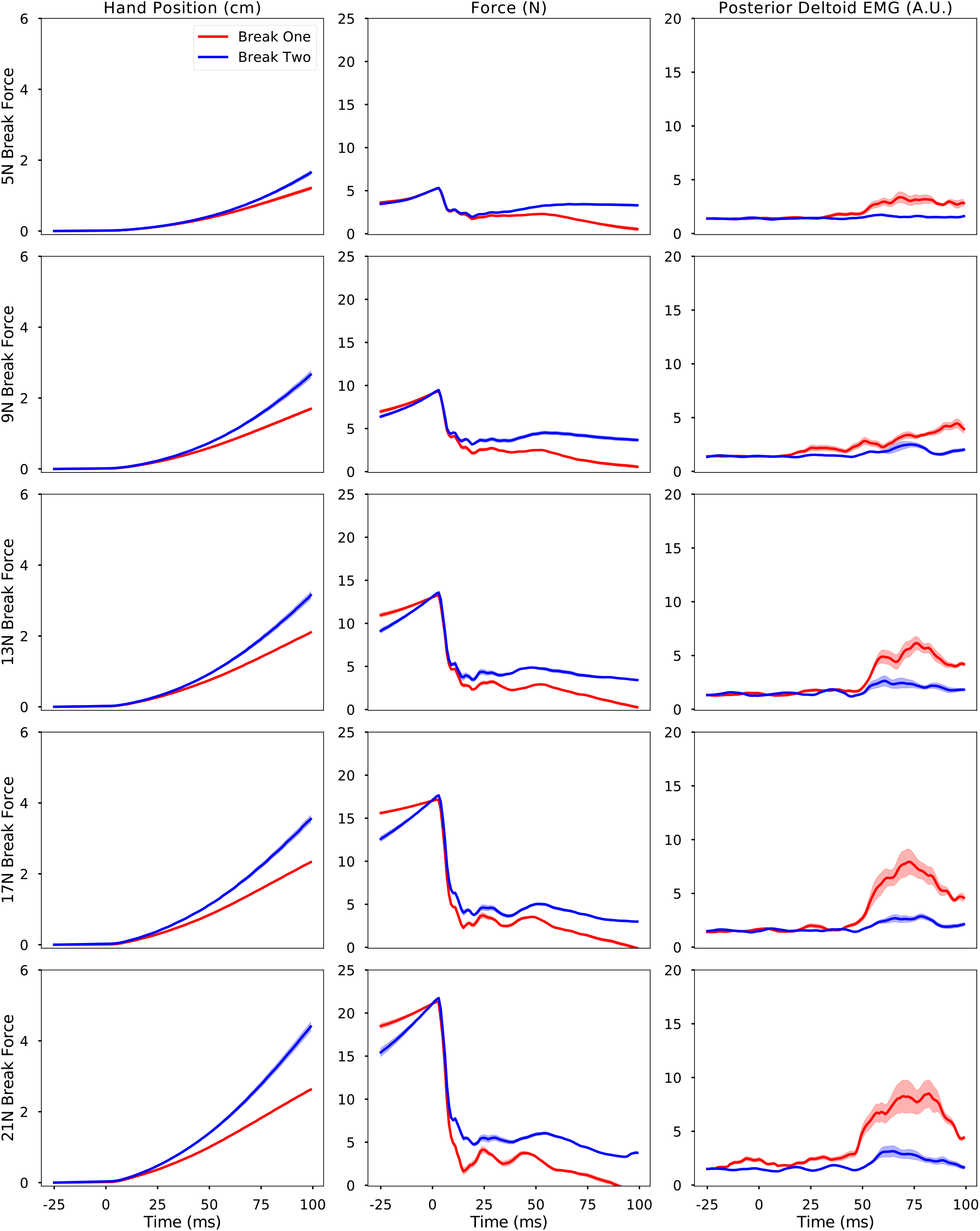
Traces of individual participant (n = 10) mean performance during Experiment 2, aligned with perturbation onset at time = 0. These data are from trials where the second wall was in the “far” position. Layout and color scheme is the same as in Figure 2.

**Figure 6.**
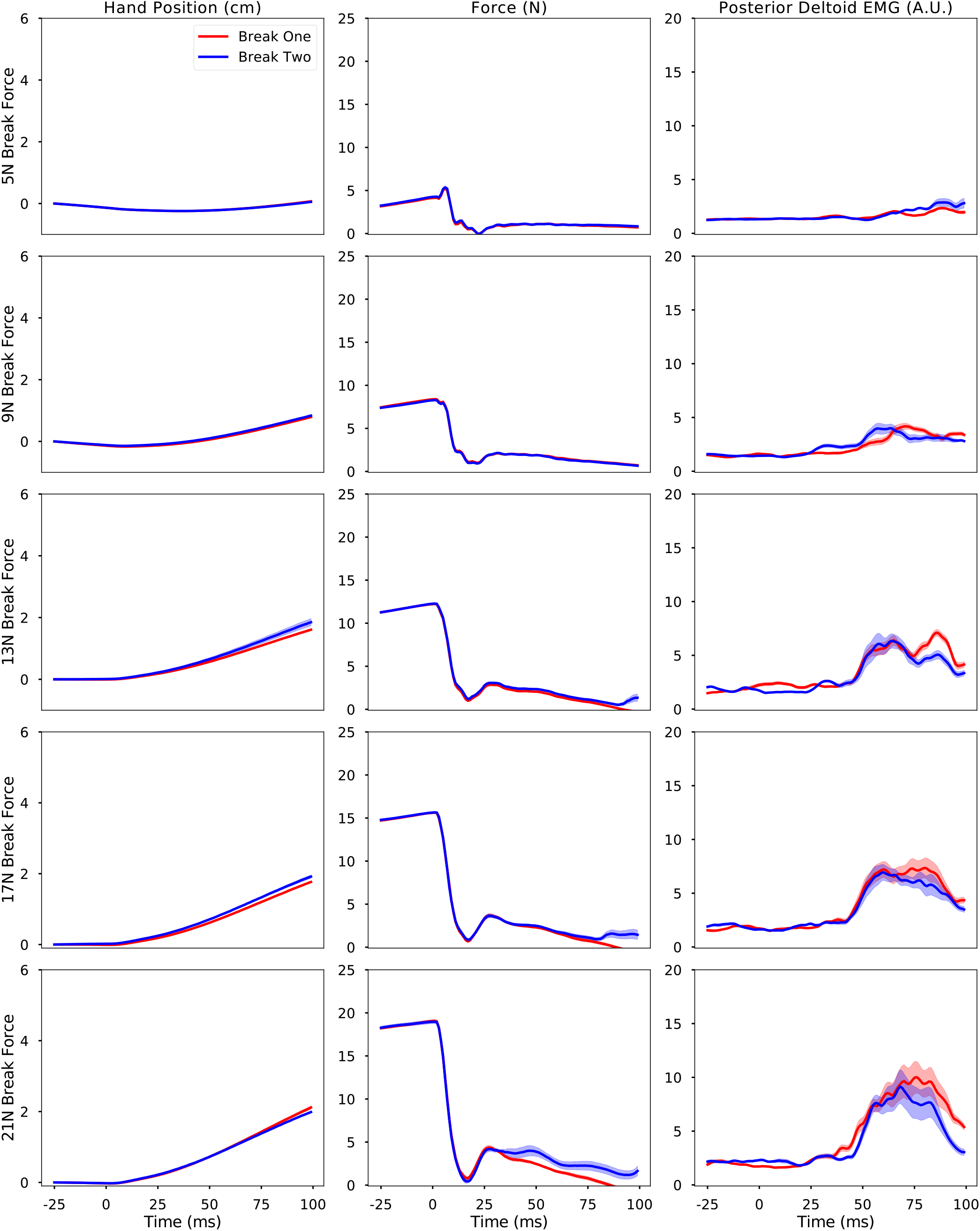
Traces of individual participant (n = 10) mean performance during Experiment 3, aligned with perturbation onset at time = 0. These data are from trials where the robot-imposed force ramp was fast. Layout and color scheme is the same as in Figures 2 and 5.

To assess whether the differences in behavior following unloading between Experiment 3 and Experiments 1 and 2 could be attributed to differences in loaded muscle state, we compared agonist and antagonist muscle activity before unloading in the Experiment 2 trials where the second wall was close to Experiment 3 trials with high force rates. This comparison was chosen because these conditions were the best-matched in terms of wall distance and force rate. This analysis did not show a robust difference between experimental conditions on the relation between break force and instruction (*Experiment × Break Force × Instruction* interaction for agonist: *F*_4,171_ = 2.33; *P* = 0.058, antagonist: *F*_4,171_ = 2.43; *P* = 0.072). Overall, both agonist and antagonist EMG were slightly higher for higher levels of break force and under the “Break One” instruction during Experiment 2 than in Experiment 3. That is, when participants produced the force to break through the first wall actively, they showed minimal difference in muscle activation compared to when they had to adjust to force imposed on them by the robot handle. The marginal p-values reflect a tendency to use slightly higher levels of co-contraction during Experiment 2 versus Experiment 3.

## Discussion

### Summary

We carried out this study to investigate upper limb responses to an ecologically common type of perturbation: one which is triggered by the agent, but is unpredictable because the triggering threshold is unknown. As a result, neither the onset nor the magnitude of the perturbation is predictable. We found that participants modulate the way they generate the force required to trigger the unloading perturbation depending on what they plan to do after unloading (Experiment 1 and 2): when participants planned to break the second wall, they increased force at a much higher rate than when they planned to arrest their movement following the unloading perturbation. Moreover, participants modulated their reflex responses to such perturbations depending on their goal following the perturbation (Experiment 1 and 2). In contrast, participants did not modulate their reflexes when they countered forces generated by the robot before the unloading event (Experiment 3).

### The role of pre-perturbation activity

Assessments of responses to perturbation tasks commonly rely on carefully controlling pre-perturbation muscle state. This control is motivated by the concept of “automatic gain-scaling” (Matthews 1986; Bedingham & Tatton 1984; Pruszynski et al 2011), in which the size-recruitment order of motor neurons causes reflex sensitivity to be proportional to the force level at which they are evoked. A key result in Experiments 1 and 2 is that, although we observed automatic gain scaling (see slopes across instructions in **Figure 3**), the task-dependent modulation was of a different origin (see different intercepts across instructions in **Figure 3**). This means that participants did not compensate simply by increasing co-contraction and relying on gain-scaling to increase the resistive force they applied following the perturbation.

### Active force production and a posture/movement neural dichotomy

Even though participants showed gain-scaling in Experiment 3 similar to their performance in Experiments 1 and 2, they did not demonstrate reliable task-dependent modulation of their reflex responses. The lack of modulation occurred despite the robot increasing force more slowly than in Experiments 1 and 2, giving participants more time to plan a perturbation response.

We speculate that this difference arises because participants are engaged fundamentally different tasks when the unloading perturbation occurs. In Experiment 3, the participants are engaged in maintaining a set hand position in response to external forces generated by the robot and the unloading perturbation acts as a cue to move their hand to a new position. In contrast, in Experiments 1 and 2, participants do not have to actively control the position of their hand before the unloading perturbation; they actively press against a virtual wall, which reactively applies the appropropriate forces to maintain hand position. As such, participants could perform Experiments 1 and 2 as movements to a known target from an unpredictable starting state.

There is evidence that posture (i.e. position maintenance) and movement rely on partially differentiated neural circuits (for exhaustive review, see Shadmehr 2017). At the behavioral level, people show minimal generalization between force fields learned during posture and reaching (Scheidt and Ghez, 2007), and dissociable deficits in trajectory and end position have been reported in stroke survivors with different lesion sites (Schaefer et al 2007). Gains on various sensory modalities are also modulated during the transition between posture and movement (Tisserand et al. 2018). In fact, even stretch reflexes change between posture and movement: Mortimer and Colleagues (1981) showed that long-latency reflexes are modified when participants prepare to make a rapid, self-initiated movement during a posture maintenance task in response to an auditory signal. Neural recordings from primary motor cortex indicate the existence of separate populations of neurons that are tuned to movement and posture tasks. For example, some monkey M1 neurons show distinct tuning to self-initiated force production, but no such tuning for bias forces counteracted before initiating that force pulse (Georgopolous et al. 1992), or distinct directional tuning at the initiation of a force pulse but not during sustained force production (Shalit et al. 2012). Similarly, during reaching, Kurtzer and Colleagues (2005) recorded separate subpopulations of neurons which were tuned to loads during posture and during movements, as well as neurons which were active during both movement and posture but changed the strength of their tuning depending on whether the monkey was currently engaged in a movement or posture task.

Participants in Experiments 1 and 2 could engage these separate neural populations during the self-generated force production by performing those tasks as a continuous movement to a position in front of the second virtual wall. If they did this, they might be able to use circuits which they could not in Experiment 3, and take advantage of those circuits for more accurate starting state estimates and/or faster access to the periphery. Alternatively, the lack of instruction-modulated reflex response in Experiment 3 could be explained by an additional switching cost of changing from a posture task to a movement task when prompted by the unloading perturbation (as proposed by Cluff & Scott, 2016).

### Our paradigm

Unloading paradigms have been used by many previous groups to study a variety of behavioral phenomena. Some studies have used unloading events triggered by an experimenter to evoke postural perturbations (Angel et al 1965, Lowrey et al 2019) or investigate muscle characteristics under do-not-intervene instructions (Archambault et al 2005). Other studies have used self-triggered unloading perturbations to investigate preparation for expected perturbations to the upper limb (Lum et al. 1992, Johansson & Westling 1988, Kennedy & Schwartz 2018) or whole body (Aruin & Latash 1995), or to trigger perturbations with unknown directions (Piscitelli et al. 2017).

Our paradigm differs from the aforementioned studies because participants self-triggered the perturbations they experienced, but the trigger thresholds were unknown. This means that participants could not accurately predict when the onset would occur or the magnitude of the perturbation. Additionally, the perturbation in our paradigm acts on the same effector as the triggering action, rather than in a relatively orthogonal manner (e.g. a pressing a button to trigger a postural perturbation as in Piscitelli et al, or opening the hand to trigger a perturbation in the other hand as in Johannson & Westling). Such perturbations are common in everyday life: for example, they occur during almost all transitions from static to dynamic friction, and every time we open a new refrigerator. Paradigms like ours can, therefore, provide an important insight into preparatory and reactive activity when people try to maintain control during this common class of unpredictable but self triggered mechanical perturbations.

Task-dependent scaling of muscle responses has been reported in previous studies where unloading perturbations were induced in the jaw muscles (Johansson et al. 2014a,b). Despite the different systems involved, the results we report are qualitatively similar; in particular, these studies again show a task-specific reflex modulation that is independent from automatic gain scaling (cf. Johansson et al. 2014a, Fig 3). In these studies, differential recruitment of compartments of the masseter muscle (which have different fiber composition) was proposed as one mechanism responsible for task-specific responses (Johansson et al 2014b). The upper limb task employs more muscles, and it is possible that complex temporal shifts in recruitment order of antagonist synergists make up some element of the response modulation we observed. Addressing this issue would require systematic coverage of many more upper-limb muscles.

Kennedy and Schwartz (2018) report preliminary findings from a task which is similar to ours in its use of a self-triggered unloading perturbation. Their primary analysis relates to modulation of co-contraction to attain various target positions following the unloading perturbations, and it is unclear whether the control strategy they report applies to our task. An important difference between that study and our own is that we completely randomized the presentation of threshold forces for unloading, while their study presented participants with the same thresholds twenty times in a row. As a result, participants learned the approximate force thresholds they had to break through, a process which likely engages different strategies as evidenced by the qualitatively different pre-unloading strategies used by their participants (see “Force” panel in **Figure 1**).

## Acknowledgements

We thank Carola Hjalten for logistical support with participant recruitment and data collection. Where possible we have tried to use free and open-source software to analyze and visualize our results. In addition to the Python and R packages previously cited, we gratefully acknowledge the individuals responsible for developing and maintaining Inkscape (inkscape.org), as well was the following components of the ScyPy ecosystem: NumPy (Oliphant 2006), Matplotlib (Hunter 2007), and Pandas (McKinney 2010).

## Contributions

S.R. drafted the manuscript; S.R., A.S.J, and J.A.P edited the manuscript; A.S.J. and J.A.P. designed the research; A.S.J. collected the data; S.R. analyzed the data; S.R. and J.A.P interpreted the findings.

